# Myonuclear Dynamics After Skeletal Muscle Surgical Injury

**DOI:** 10.64898/2026.05.12.724630

**Authors:** Micah Goeke, Nathan Serrano, Pieter Jan Koopmans, Kevin A. Murach

## Abstract

A hallmark of damaged skeletal muscle fibers is displaced myonuclei that are no longer peripherally positioned. Displaced myonuclei are dogmatically thought to be derived exclusively from muscle stem cell (satellite cell) fusion. Using a surgical resection muscle injury model and *in vivo* recombination-independent resident myonuclear labeling, we detail the prevalence, time course, and origin of displaced myonuclei in response to a non-chemically-mediated muscle trauma. We found that: 1) non-satellite cell-derived (resident) displaced myonuclei emerge seven days after surgical injury in similar proportion to exogenous (satellite cell-derived) displaced myonuclei in intact muscle fibers, with a biased prevalence in myosin heavy chain IIB muscle fibers, 2) muscle fibers with multiple (≥2) displaced resident myonuclei was an unexpected but noteworthy feature of muscle fibers 7 days after injury, 3) embryonic myosin-expressing fibers at seven days post-surgery expectedly contain predominantly satellite-cell derived displaced myonuclei, but a subset have displaced resident myonuclei, and 4) satellite cell numbers in intact muscle do not increase until 7 days post-surgery. These data may help inform whether to target satellite cell-initiated processes, myonuclear-initiated processes, or both to facilitate muscle fiber injury repair. This information could lead to more effective therapeutic strategies for treating muscle trauma.

## Main text

The majority of contemporary knowledge on how skeletal muscle responds to severe trauma has been gathered using models involving myotoxic chemical injections (e.g. barium chloride, cardiotoxin, notexin, etc.) (1, 2), freeze/thermal (2, 3), crush (4), or ischemia-reperfusion injury (5), volumetric muscle loss (6), or eccentric contraction damage (7, 8). In reality, for humans, the most common form of severe skeletal muscle injury likely occurs during surgery. Muscle is incised to access underlying anatomy or excised (and often discarded) for various reasons. A few classic light and electron microscopy studies showed observations using laceration-type injuries (9, 10), but in general, detailed studies on how skeletal muscle responds to surgical injury at the cellular and organelle level are generally scarce (11, 12). Furthermore, the role of muscle stem cells (satellite cells) has overwhelmingly been the focus of muscle injury repair research. This focus is largely due to the indispensable function of satellite cells in skeletal muscle regeneration (13-16).

A hallmark of degenerated then regenerated skeletal muscle fibers is displaced or centralized myonuclei. These displaced myonuclei are in contrast to the typical peripheral positioning of myonuclei and are attributed to the activities of satellite cells (17). In intact adult myofibers that experience damage, however, there is recent evidence that displaced myonuclei can be the result of resident (non-satellite cell-derived) myonuclear translocation (18-20). Resident peripherally-located myonuclei can migrate for the purpose of promoting cell-autonomous muscle fiber repair processes (20, 21). Understanding the contributions of resident myonuclei to the injury response is therefore a burgeoning area of interest (21, 22). Whether to target satellite cell processes, myonuclear processes, or both to promote muscle fiber injury repair could lead to more effective and/or combinatorial therapeutic strategies for treating muscle trauma.

The purpose of this investigation was to understand the timing and contributions of resident versus satellite-cell derived myonuclei in the context of surgical muscle injury. Our readout was a classic hallmark of muscle damage repair: displaced myonuclei. Two unique approaches were used to accomplish our goal: 1) a doxycycline-inducible recombination-independent model of fluorescent myonuclear labelling (called HSA-GFP) that allows for high-confidence discrimination between resident myonuclei and those acquired from an exogenous source (such as satellite cells) in adult skeletal muscle (18, 19, 23-25), and 2) a surgical resection model where the behavior of intact skeletal muscle fibers – and therefore resident myonuclei – can be studied while minimizing complete muscle degeneration. We also delivered EdU (5-ethynyl-2’-deoxyuridine) to assess DNA synthesis and satellite cell contributions to muscle fibers and quantified PAX7+ satellite cell abundance.

### Study design

Adult HSA-GFP mice (∼10 month old) were given 0.5 mg/ml doxycycline and 2% sucrose for five days in drinking water to fluorescently label resident myonuclei. After a 14-day washout period, mice underwent bilateral muscle resection surgery. The surgery involved removal of the lower ∼1/3 of the gastrocnemius/soleus complex and achilles tendon (Figure 1A). Mice returned to normal ambulation (relying on the plantaris muscle for plantarflexion) within ∼24 hours (18). Resected muscles were collected three days (n=4; M/F=2/2) or seven days after surgery (n=4; M/F=2/2), and sham operated mice (no resection and time-matched to the 7-day cohort, n=4; M/F=2/2) were controls. At the time of surgery, mice were injected with an intraperitoneal “priming dose” of EdU (∼2 mg in water) then given 0.5 mg/ml EdU (Biosynth, NE08701) with 2% sucrose in drinking water that was shielded from light and refreshed every other day; this approach results in appreciable DNA labeling in our hands (18). Resected muscles were prepared for histology according to our standard procedures (18) and analyzed for displaced myonuclei: resident GFP+/DAPI+ or exogenous GFP-/DAPI+ and not abutting the dystrophin border (called GFP+ or GFP- from here onward), quantified according to myosin heavy chain 2B expression (MyHC IIB+ or IIB-, with IIB+ being pure IIB and/or IIB/IIX, and IIB- likely being IIA and/or IIX fibers) or embryonic myosin heavy chain fiber type expression (eMyHC+ or eMyHC-). The muscles were analyzed a few millimeters away from the injury site. Displaced myonuclei were also quantified as EdU+ or EdU-. Satellite cells per fiber were identified as PAX7+/DAPI+ and within the muscle fiber laminin border (26, 27).

**Figure 1.**
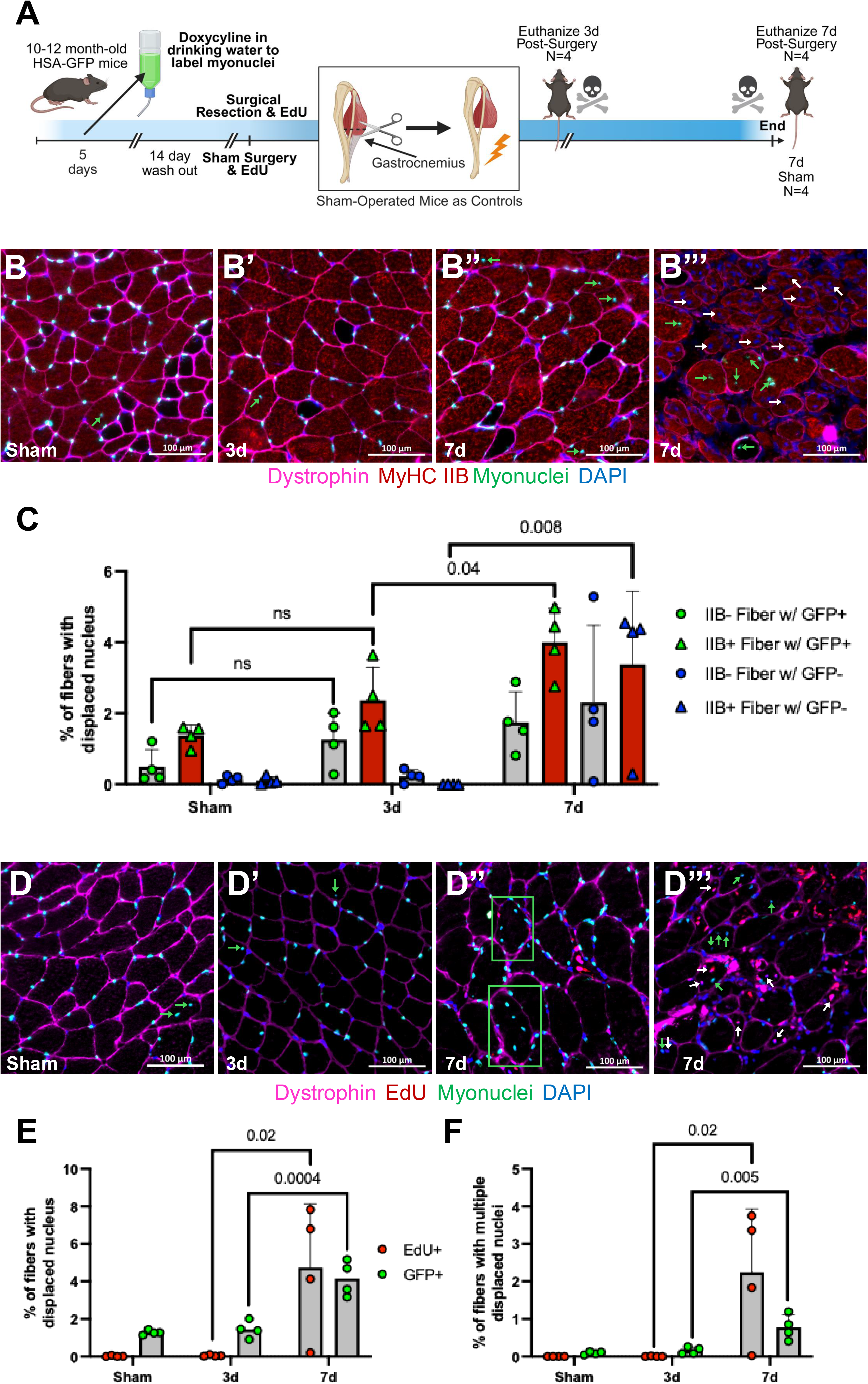
**A)** Study design where 10-month-old HSA-GFP mice (M/F=6/6) were treated with doxycycline to label myonuclei for 5 days followed by a 14-day wash-out. The distal ∼1/3 of the gastrocnemius-soleus complex was excised and mice were treated with 5-ethynyl-2’-deoxyuridine (EdU) in drinking water to assess DNA synthesis following surgical resection injury. Resected gastrocnemius muscles were collected at three days (n=4; M/F=2/2) and seven days (n=4; M/F=2/2) post-surgery, with sham controls collected at 7 days (N=4; M/F=2/2). **B**-**B’’’)** show representative images of the gastrocnemius muscle for fiber type-specific quantification of displaced myonuclei. Images show dystrophin (pink), myosin heavy chain IIB (MyHC IIB, red), GFP-labeled resident myonuclei (green), and DAPI nuclei (blue). Green arrows = GFP+ resident myonuclei, white arrows = GFP- non-resident myonuclei. **C**. Quantification of relative number of IIB+ or IIB- fibers containing one or more displaced myonuclei. Data are presented as mean ± SD and is reported relative to each respective fiber type. **D**-**D’’’)** Representative images of the gastrocnemius muscle for DNA synthesis analysis stained for dystrophin (pink), DAPI (blue), and 5-ethynyl-2’-deoxyuridine (EdU, red), with GFP+ resident myonuclei in green. Green arrows = GFP+/EdU- resident myonuclei, white arrows = GFP-/EdU+ exogenous myonuclei. **E)** Quantification of relative number of fibers containing multiple GFP+/EdU- or GFP-/EdU+ myonuclei. **F)** Quantification of relative number of fibers containing multiple GFP+ or EdU+ nuclei is shown in Data are presented as mean ± SD. Significance was established *a priori* at *p*<0.05 and assessed *via* one-way ANOVA.

### Three-day post-surgical myonuclear response

Three days after surgical resection, GFP+ (resident) displaced myonuclei were not significantly more abundant relative to sham in MyHC IIB- or IIB+ muscle fibers (Figures 1B and 1C); however, IIB+ fibers were beginning to feature modestly more displaced resident myonuclei (1.4% vs 2.4%, *p* = 0.23). There was greater variability in displaced resident myonuclei in both fiber types with surgery relative to sham; peraps a larger sample size may had revealed significance. GFP- (presumably satellite cell-derived) displaced myonuclei were not more abundant in MyHC IIB+ or IIB- muscle fibers three days after surgical resection (∼0% in both fiber types). In response to mechanical overload (MOV) of the plantaris muscle, IIB- fibers had significantly more displaced resident myonuclei versus sham after three days in our hands (18). Mechanical tension may, in part, drive early myonuclear translocation in IIB- muscle fibers, whereas damage alone may not. Relative to the plantaris, though, IIB- (IIA and/or IIX) fibers are less prevalent in the gastrocnemius. This relative scarcity may also contribute to the conclusions from our analysis.

### Seven-day post-surgical myonuclear response

Seven days after surgical resection, IIB- fibers did not have significantly more GFP+ of GFP- displaced myonuclei; however, there was again more variability relative to sham (Figure 2C). IIB+ muscle fibers had more displaced GFP+ and GFP- displaced myonuclei seven days after injury relative to sham (IIB+/GFP+: *p* = 0.002; IIB+/GFP-: *p* = 0.009), and more versus three days after surgery (IIB+/GFP+: *p* = 0.04; IIB+/GFP-: *p* = 0.008) (Figures 1B and 1C). In total, IIB+ fibers had ∼4% of fibers with GFP+ and ∼3.5% GFP- displaced myonuclei seven days after injury (∼7.5% of total fibers). This overall prevalence exceeds what is found after seven days of MOV (<4% of total fibers with displaced myonuclei) (18). There were some regional differences where intact myofibers predominantly contained GFP+ displaced myonuclei whereas regions undergoing degeneration (smaller fibers with large interstitial spacing) were dominated by GFP- displaced nuclei (Figures 1B’’ and 1B’’’, respectively). One sample had an unexpected phenotype where GFP+ displaced myonuclei were prevalent seven days after injury but GFP- were not (Figure 1C). Our analysis of this muscle may had been more distant from the injury site versus the others, or perhaps a portion of resected muscle had actually remained intact in this sample. Nevertheless, we elected to retain this sample in the analysis. The timing of events – where GFP+ displaced myonuclei may begin emerging early after injury and become more prevalent over time – agrees with classic work where myofiber membrane sealing (cell-autonomous process) precedes satellite cell fusion (non-cell-autonomous process) after injury (28), and reinforces speculation where myonuclear migration explains early myonuclear accumulation at the damaged ends of intact myofibers (9, 28).

**Figure 2.**
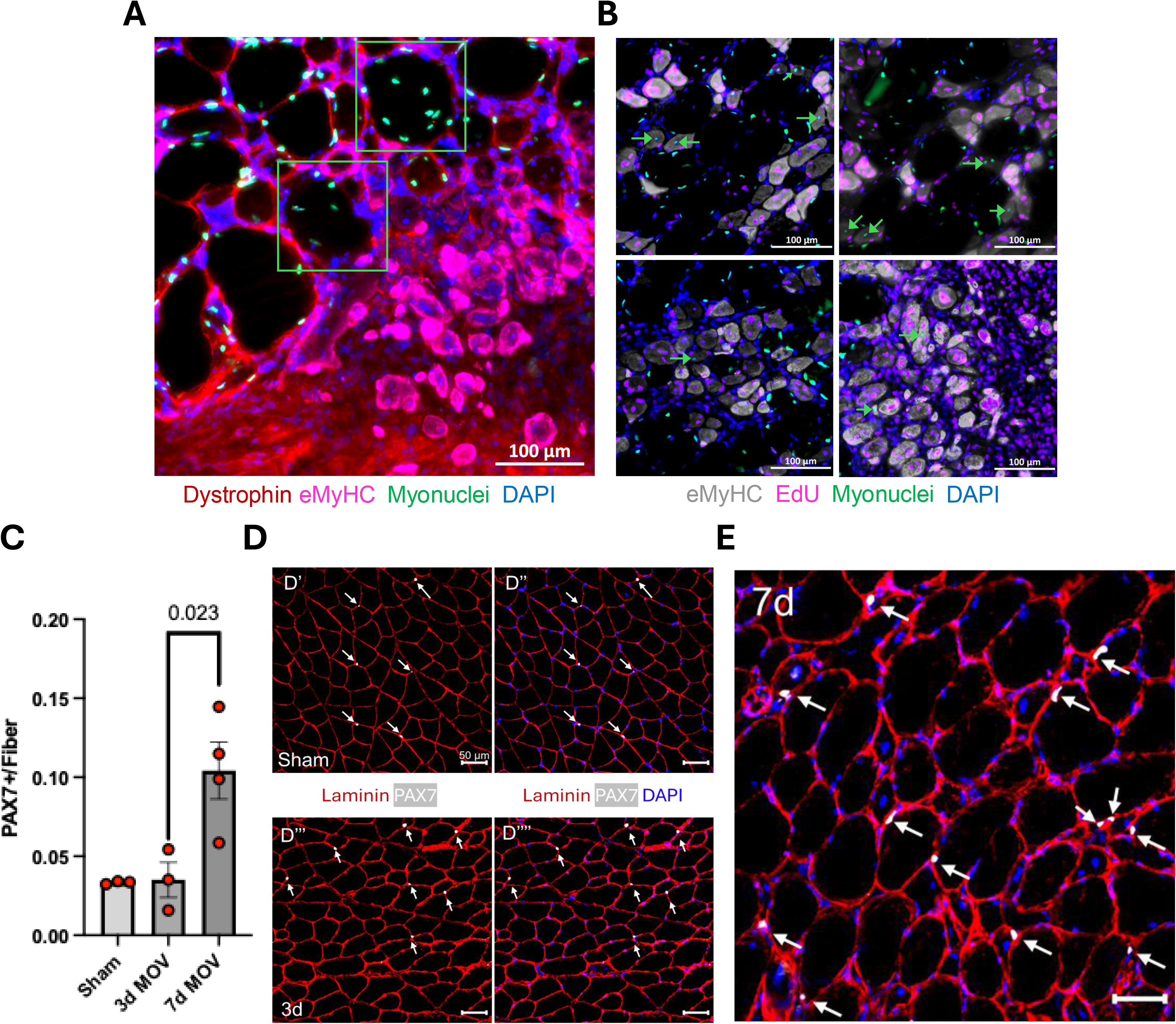
**A)** Z-stack image from 30 μm section of seven day gastrocnemius showing regional eMyHC abundance and prevalence of several displaced resident myonuclei within a single fiber. **B)** Representative images of emybronic myosin positive fibers with displaced resident myonuclei. Green arrows = GFP+ resident myonuclei. **C**. Quantification of number of satellite cells at each timepoint. Data are presented as mean ± SD and are reported relative to the number of fibers. **D)** Representative images of satellite cells in the gastrocnemius muscle at sham (D and D’) and three day timepoints (D’’ and D’’’). Laminin (red), PAX7 (white), and DAPI (nuclei, blue). **E)** Representative image of satellite cells (white arrow) in the gastrocnemius muscle at seven day timepoint. Scale bar is 50 μm in panels D and E.

### DNA synthesis analysis of displaced myonuclei three and seven days after surgery

Nearly all displaced GFP+ myonuclei in injured muscle were EdU-, and displaced GFP- were EdU+; the latter were most likely derived from satellite cells that had proliferated and fused to the muscle fiber. Consistent with the GFP+/GFP- analysis presented above, the abundance of displaced GFP+ and EdU+ myonuclei in intact muscle fibers was not significantly higher until seven days after surgical injury (EdU+: 3 day vs 7 day: *p* = 0.02; GFP+: 3 day vs 7 day: *p* = 0.0004) (Figures 1D-D’’’ and 1E). A unique and striking feature of surgical injury compared to mechanical overload (18) with respect to displaced myonuclei was the relative preponderance of muscle fibers with multiple (≥2) displaced myonuclei after surgery (see Figure 1D’’-D’’’). There were more individual fibers with multiple displaced myonuclei that were EdU+ (likely regenerating, up to 120 fibers per muscle cross-section, ∼2.5% of all fibers); however, up to 30 fibers per muscle (<1% of all fibers) had multiple GFP+ displaced resident myonuclei (likely damaged but intact fibers) seven days after injury (EdU+: 3 day vs 7 day: *p* = 0.02; GFP+: 3 day vs 7 day: *p* = 0.005) (Figures 1D’’-D’’’ and 1F). We obtained thicker muscle sections (30 µm) to better visualize these fibers and myonuclei (Figure 2A). Unlike the organized and linear “myonuclear chains” that often occur during regeneration (17, 29), it appears that these GFP+ myonuclei were scattered throughout the inside of the muscle fiber. Future studies may use isolated individual muscle fibers and three-dimensional imaging to more clearly characterize these fibers and myonuclei. Few fibers had multiple displaced GFP+ myonuclei in the sham condition (Figure 1F). Our observations provide evidence that resident myonuclei can mobilize *en masse* in response to surgical damage in intact adult muscle fibers.

### eMyHC+ muscle fibers with GFP+ displaced myonuclei

Regenerating muscle fibers expressing eMyHC would be expected to emerge in response to severe injury. Since regeneration and eMyHC expression is driven by satellite cells (13), it is intuitive that the majority of displaced myonuclei in eMyHC+ fibers would be the result of satellite cell fusion (GFP- and/or EdU+). However, we previously reported that GFP+ displaced resident myonuclei can appear in eMyHC+ fibers in adult skeletal muscle in response to stress (18, 19). We therefore sought to evaluate eMyHC+ muscle fibers with GFP+ displaced myonuclei. These fibers were rare, comprising a small proportion of all eMyHC+ fibers (Figure 2B). Nevertheless, resident displaced myonuclei in eMyHC+ fibers may be an unrecognized but potentially meaningful feature of muscle fiber degeneration/regeneration (19). We also anecdotally observed that the further away we moved from the injury site, the less eMyHC was apparent. This suggests that eMyHC localizes to the site of injury where the most satellite cell fusion is likely occurring.

### Satellite cell abundance after surgical resection injury

With our surgical resection approach, the achilles tendon is excised along with the muscle which leads to the removal of muscle loading on the resected muscle. Surgical resection is therefore a damage model simultaneous with an atrophy stimulus (i.e. deloading and possible denervation). How satellite cells react to such a stimulus is not well-described. Muscle damage typically causes satellite cell proliferation (8) whereas deloading either does not affect satellite cell number (30) or reduces it (31, 32). We therefore evaluated the abundance of PAX7+ satellite cells to further characterize the resection model in intact muscle fibers where clear laminin borders were discernible. At 3 days, satellite cell abundance in intact regions of muscle was not changed (Figure 2C and 2D-D’), but at 7 days satellite cells were ∼3-fold more abundant (sham vs 7d, *p* = 0.021; 3d vs 7d, *p* = 0.023) (Figures 2D and E). In some regions undergoing active regeneration, satellite cells were very abundant but could not be easily quantified due to disrupted laminin architecture.

## Conclusions

We report that displaced resident myonuclei appear in response to unaccustomed physical activity in the soleus muscle (19) and mechanical overload of the plantaris muscle (18), but most prominently with surgical resection in the gastrocnemius of adult muscle shown here. In the current study, we cannot confidently say whether every fiber in cross-section was directly damaged, in part due to the pennation angles of the medial and lateral gastrocnemius muscles. Nevertheless, in non-regenerating (eMyHC-) intact muscle fibers, displaced resident myonuclei were a prominent feature that is directly related to damage or indirectly related to inflammation, denervation, and/or deloading. In our hands, resident myonuclear movement is a conserved feature across muscle types and stressors. Perhaps in the early phases of injury, targeting myonuclear motility processes may help improve intrinsic muscle fiber healing, whereas later phases may benefit from targeting satellite cells.

## Methods

### Ethical approval

This study was approved by the Institutional Animal Care and Use Committee at the University of Arkansas and all procedures were performed in accordance with the National Institutes of Health (NIH) *Guide for the Care and Use of Laboratory Animals*. All mice were housed at 21-23°C under pathogen-free conditions with five or fewer mice per cage. Mice had free access to food and water.

### Mouse study methods

#### Experimental Design

Adult 10-month-old HSA-GFP mice (N=12; M/F=6/6) were treated with 0.5mg/ml doxycycline and 2% sucrose for 5 days in their drinking water to fluorescently label resident myonuclei. Following a 14 -day washout period, mice underwent bilateral muscle resection surgery. Under an isoflurane anesthetic, the Achilles tendon was cut, and the distal 1/3-1/2 of the gastrocnemius-soleus complex was excised. Mice returned to normal ambulation immediately. Resected gastrocnemius muscles were collected at three days (n=4; M/F=2/2) and 7 days (n=4; M/F=2/2) post-surgery. Sham-operated mice (with no resection) were time-matched to the seven-day cohort to serve as controls (n=4; M/F=2/2). To assess DNA synthesis and track exogenous cell fusion, mice were administered an intraperitoneal “priming dose” of 5-ethynyl-2’-deoxyuridine (EdU, ∼2 mg in water) at the time of surgery, then give 0.5 mg/ml EdU (Biosynth, NE08701) with 2% sucrose in drinking water, refreshed every other day.

#### Histology and Immunohistochemistry

At euthanasia, gastrocnemius muscles were collected and embedded in optimal cutting temperature (OCT) compound, snap-frozen in liquid-nitrogen chilled isopentane, and sectioned at 7 µm or 30 µm (specified in figure legends) using an Epredia Cryostar NX50 Cryostat. For fiber type-specific assessment of displaced nuclei (no EdU), sections were allowed to air dry for 1 hour and encircled using a hydrophobic-barrier PAP Pen (ImmEdge pen, cat. no. H-4000, Vector Laboratories, Burlingame, CA, USA). Tissue sections were incubated with a primary antibody cocktail of 1:100 dystrophin rabbit anti-mouse IgG (ab15277, abcam, Cambridge, UK) and 1:100 myosin heavy chain (MyHC) IIb IgM (BF-F3, DSHB) for > 4 h. Tissues were washed for three rounds of three minutes in 1X PBS, then incubated in a secondary antibody cocktail of 1:200 donkey anti-rabbit IgG (Alexa Fluor 647, AB_2340626) and 1:200 goat anti-mouse IgM. Tissues were again washed for three rounds of five minutes in 1X PBS, and DAPI was applied post-wash for 10 minutes. Tissues were subsequently washed for two rounds of two minutes in 1X PBS, mounted with a 50% glycerol medium and whole section coverslips, then imaged.

For non-fiber-type-specific analysis of displaced nuclei, sections were allowed to air dry for one hour and encircled with a PAP pen as previously described. Tissue sections were incubated in a 1:100 dystrophin IgG primary antibody solution for > 4 hours followed by three rounds of three minute washes in 1x PBS. Tissues were then incubated for 90 min in a secondary antibody solution of 1:200 donkey anti-rabbit IgG (Alexa Fluor 647, magenta). Tissues were again washed for three rounds of five minutes in 1X PBS. Tissues were then post-fixed in 4% paraformaldehyde (PFA) for 15 minutes and washed for three rounds of five minutes. EdU detection was then performed with a single CLICK-iT detection buffer for 90 min containing 100 mM Tris, 5 mM Copper Sulfate, 100 mM ascorbic acid, and TAMRA-Azide (Vector, CCT-AZ109-1). Tissues were washed for three rounds of five minutes in 1X PBS. DAPI was then applied for 10 minutes, and tissues were washed in 1x PBS for two rounds of three minutes. Tissues were mounted as previously described and imaged. Displaced nuclei were defined as myonuclei not abutting the dystrophin boundary (sarcolemma) and were manually assessed in Zen software.

For embryonic myosin heavy chain (eMyHC) staining, tissue sections were allowed to air dry for 1 hour and encircled with a PAP pen as previously described. Tissues were blocked with 1% BSA and 10% normal horse serum in 1X PBS for 30 minutes at room temperature and washed for three rounds of three minutes in 1X PBS. Tissues were incubated in a 1:100 eMyHC IgG1 (F.1652, DSHB) primary antibody solution for > 4 hours. Tissues were then washed for three rounds of five minutes in 1X PBS. Tissues were incubated for 90 minutes in a secondary antibody solution of 1:200 goat anti-mouse IgG1 (Alexa Fluor 647, Magenta). Tissues were washed for three rounds of five minutes in 1X PBS, and EdU detection was performed as previously described.

For PAX7+ nuclei, tissue sections were fixed with 4% PFA for seven minutes and washed for three rounds of three minutes in 1X PBS, treated with 3% hydrogen peroxide to block endogenous peroxides for seven minutes, followed by an epitope retrieval step in sodium citrate (2.94g/L pH 6.8) at 92°C for 11 minutes and allowed to cool to room temperature and washed for three rounds of three minutes in PBS. Sections were then blocked with mouse-on-mouse IgG blocking reagent (vector, #MKB-2213) for 45 minutes followed by three rounds of PBS washes, and then with 1% TSA with 0.1% Triton at room temperature for one hour. Anti-PAX7 (ab_528428, DSHB) and laminin (Sigma L9393) primary antibodies were then applied overnight at 4°C. Following biotinylated secondary antibody (1:1000) treatment in 1% TSA (Jackson Immunoresearch, 115-065-205) for 70 minutes and three rounds of three minute 1X PBS washes, tissues were then treated with SA-HRP (1:500; Thermo Fisher, S-911) and laminin (anti-rabbit AF647; 1:200) secondaries for one hour at room temperature before another round of PBS washes and applying the TSA AF594 amplification for PAX7 for 15 minutes. Nuclei were stained with Prolong Diamond anti-fade mountant with DAPI (Invitrogen, P36962).

#### Image capture and analysis

All images were captured using an upright fluorescent microscope at 20X magnification (Zeiss AxioImager M2, Oberkochen, Germany). Whole muscle cross-sections were imaged using the mosaic function in Zeiss Zen 3.8 for Microsoft. Manual quantification of displaced resident myonuclei (GFP+, DAPI+), proliferating non-myonuclei (GFP-, DAPI+, EdU+), and fibers with displaced nuclei were performed using Zen software. Muscle fiber counts on whole muscle cross-sections were assessed in MyoVision semi-automated analysis software^32^.

#### Statistics

Displaced nuclei counts were analyzed using GraphPad statistical software (Prism, version 11.0.0 for Windows). For all analyses, one-way ANOVAs were performed independently on relative displaced myonuclear counts according to the individual populations over time (Sham, 3d, 7d): IIB+/GFP+, IIB+/GFP-, IIB-/GFP+ and IIB- /GFP- nuclei. For proliferative status populations were: GFP+/EdU-, GFP-/EdU+, and fibers containing multiple (≥2) displaced nuclei. Tukey’s post-hoc corrections were applied to all ANOVAs. Statistical significance was set at *p* ≤ 0.05. All figures were created using GraphPad Prism.

## Authors’ Contributions

NS, PJK, and KAM conceived of the study analysis approaches. MG, NS, and PJK analyzed data. KAM wrote the manuscript with input from MG, NS, and PJK. MG, NS, and PJK generated figures. MG, NS, PJK, and KAM managed and/or performed experiments. KAM provided resources, oversight, and/or intellectual contributions. All authors provided feedback and final approval of the manuscript.

## References

1. Tierney, M. T., and Sacco, A. (2016) Inducing and evaluating skeletal muscle injury by notexin and barium chloride. In Skeletal Muscle Regeneration in the Mouse: Methods and Protocols pp. 53–60, Springer

2. Hardy, D., Besnard, A., Latil, M., Jouvion, G., Briand, D., Thépenier, C., Pascal, Q., Guguin, A., Gayraud-Morel, B., and Cavaillon, J.-M. (2016) Comparative study of injury models for studying muscle regeneration in mice. PloS One 11, e0147198

3. Brightwell, C. R., Hanson, M. E., El Ayadi, A., Prasai, A., Wang, Y., Finnerty, C. C., and Fry, C. S. (2020) Thermal injury initiates pervasive fibrogenesis in skeletal muscle. Am J Physiol Cell Physiol 319, C277–C287

4. Utvåg, S. E., Grundnes, O., Rindal, D. B., and Reikerås, O. (2003) Influence of extensive muscle injury on fracture healing in rat tibia. J Ortho Trauma 17, 430–435

5. Sicherer, S. T., Venkatarama, R. S., and Grasman, J. M. (2020) Recent trends in injury models to study skeletal muscle regeneration and repair. Bioengineering 7, 76

6. Pollot, B. E., and Corona, B. T. (2016) Volumetric muscle loss. In Skeletal Muscle Regeneration in the Mouse: Methods and Protocols pp. 19–31, Springer

7. Højfeldt, G., Hoegsbjerg, C., von Keudell, A. G., and Mackey, A. L. (2025) The repair capacity spectrum of human skeletal muscle injury from sports to surgical trauma settings. J Physiol 603, 7441–7454

8. Mackey, A. L., and Kjaer, M. (2017) The breaking and making of healthy adult human skeletal muscle in vivo. Skelet Muscle 7, 24

9. Ali, M. (1979) Myotube formation in skeletal muscle regeneration. J Anat 128, 553–562

10. Hall-Craggs, E. (1974) The regeneration of skeletal muscle fibres per continuum. J Anat 117, 171

11. Garrett Jr, W. E., Seaber, A. V., Boswick, J., Urbaniak, J. R., and Goldner, J. L. (1984) Recovery of skeletal muscle after laceration and repair. J Hand Surg 9, 683–692

12. Chan, Y.-S., Li, Y., Foster, W., Horaguchi, T., Somogyi, G., Fu, F. H., and Huard, J. (2003) Antifibrotic effects of suramin in injured skeletal muscle after laceration. J Appl Physiol 95, 771–780

13. McCarthy, J. J., Mula, J., Miyazaki, M., Erfani, R., Garrison, K., Farooqui, A. B., Srikuea, R., Lawson, B. A., Grimes, B., Keller, C., Van Zant, G., Campbell, K. S., Esser, K. A., Dupont-Versteegden, E. E., and Peterson, C. A. (2011) Effective fiber hypertrophy in satellite cell-depleted skeletal muscle. Development 138, 3657–3666

14. Lepper, C., Partridge, T. A., and Fan, C.-M. (2011) An absolute requirement for Pax7-positive satellite cells in acute injury-induced skeletal muscle regeneration. Development 138, 3639–3646

15. Murphy, M. M., Lawson, J. A., Mathew, S. J., Hutcheson, D. A., and Kardon, G. (2011) Satellite cells, connective tissue fibroblasts and their interactions are crucial for muscle regeneration. Development 138, 3625–3637

16. Sambasivan, R., Yao, R., Kissenpfennig, A., Van Wittenberghe, L., Paldi, A., Gayraud-Morel, B., Guenou, H., Malissen, B., Tajbakhsh, S., and Galy, A. (2011) Pax7-expressing satellite cells are indispensable for adult skeletal muscle regeneration. Development 138, 3647–3656

17. Pizza, F. X., and Buckley, K. H. (2023) Regenerating myofibers after an acute muscle injury: What do we really know about them? Int J Mol Sci 24, 12545

18. Serrano, N., Koopmans, P. J., and Murach, K. A. (2025) Displaced myonuclei are attributable to both resident myonuclear migration and stem cell fusion during mechanical loading in adult skeletal muscle. Skelet Muscle, 16(4).

19. Murach, K. A., Mobley, C. B., Zdunek, C. J., Frick, K. K., Jones, S. R., McCarthy, J. J., Peterson, C. A., and Dungan, C. M. (2020) Muscle memory: myonuclear accretion, maintenance, morphology, and miRNA levels with training and detraining in adult mice. J Cachex Sarc Muscle 11, 1705–1722.

20. Roman, W., Pinheiro, H., Pimentel, M. R., Segalés, J., Oliveira, L. M., García-Domínguez, E., Gómez-Cabrera, M. C., Serrano, A. L., Gomes, E. R., and Muñoz-Cánoves, P. (2021) Muscle repair after physiological damage relies on nuclear migration for cellular reconstruction. Science 374, 355–359

21. Roman, W., and Muñoz-Cánoves, P. (2022) Muscle is a stage, and cells and factors are merely players. Trend Cell Biol 32, P835–840.

22. Bagley, J. R., Denes, L. T., McCarthy, J. J., Wang, E. T., and Murach, K. A. (2023) The myonuclear domain in adult skeletal muscle fibres: past, present and future. J Physiol 601, 723–741

23. Murach, K. A., Dungan, C. M., von Walden, F., and Wen, Y. (2021) Epigenetic evidence for distinct contributions of resident and acquired myonuclei during long-rerm exercise adaptation using timed in vivo myonuclear labeling. Am J Physiol Cell Physiol 32, C86–C93

24. Wen, Y., Dungan, C. M., Mobley, C. B., Valentino, T., von Walden, F., and Murach, K. A. (2021) Nucleus type-specific DNA methylomics reveals epigenetic “memory” of prior adaptation in skeletal muscle. Function, zqab038

25. Koopmans, P. J., Jones III, R. G., Cabrera, A. R., Morena, F., Greene, N. P., McCarthy, J. J., Ismaeel, A., Wen, Y., and Murach, K. A. (2026) The age-dependent resident myonuclear multi-omic response to an acute skeletal muscle hypertrophic stimulus in mice. Adv Sci, e21633

26. Murach, K. A., Peck, B. D., Policastro, R. A., Vechetti, I. J., Van Pelt, D. W., Dungan, C. M., Denes, L. T., Fu, X., Brightwell, C. R., and Zentner, G. E. (2021) Early satellite cell communication creates a permissive environment for long-term muscle growth. iScience 24, 102372

27. Murach, K. A., Vechetti Jr, I. J., Van Pelt, D. W., Crow, S. E., Dungan, C. M., Figueiredo, V. C., Kosmac, K., Fu, X., Richards, C. I., Fry, C. S., McCarthy, J. J., and Peterson, C. A. (2020) Fusion-independent satellite cell communication to muscle fibers during load-induced hypertrophy. Function 1, zqaa009

28. Robertson, T., Papadimitriou, J., and Grounds, M. (1993) Fusion of myogenic cells to the newly sealed region of damaged myofibres in skeletal muscle regeneration. Neuropath Appl Neurobio 19, 350–358

29. Wada, K. I., Katsuta, S., and Soya, H. (2008) Formation process and fate of the nuclear chain after injury in regenerated myofiber. Anat Rec 291, 122-128

30. Jackson, J. R., Mula, J., Kirby, T. J., Fry, C. S., Lee, J. D., Ubele, M. F., Campbell, K. S., McCarthy, J. J., Peterson, C. A., and Dupont-Versteegden, E. E. (2012) Satellite cell depletion does not inhibit adult skeletal muscle regrowth following unloading-induced atrophy. Am J Physiol Cell Physiol 303, C854–861

31. Mitchell, P. O., and Pavlath, G. K. (2004) Skeletal muscle atrophy leads to loss and dysfunction of muscle precursor cells. Am J Physiol Cell Physiol 287, C1753–C1762

32. Arentson-Lantz, E. J., English, K. L., Paddon-Jones, D., and Fry, C. S. (2016) Fourteen days of bed rest induces a decline in satellite cell content and robust atrophy of skeletal muscle fibers in middle-aged adults. J Appl Physiol 120, 965–975

